# Marine particle microbiomes during a spring diatom bloom contain active sulfate-reducing bacteria

**DOI:** 10.1101/2022.05.31.494182

**Authors:** R. Siebers, D. Schultz, M. S. Farza, A. Brauer, D. Zühlke, P. A Mücke, F. Wang, J. Bernhardt, H. Teeling, D. Becher, K. Riedel, I. V. Kirstein, K. H. Wiltshire, K.J. Hoff, T. Schweder, T. Urich, M. M. Bengtsson

## Abstract

Phytoplankton blooms fuel marine food webs with labile dissolved carbon, but also lead to the formation of particulate organic matter composed of living and dead algal cells. These particles contribute to carbon sequestration, yet are also sites of intense algal-bacterial interactions and provide diverse niches for microbes to thrive. We analyzed 16S and 18S ribosomal RNA gene amplicon sequences obtained from 51 time points and metaproteomes from 3 time points during a spring phytoplankton bloom in the North Sea. Particulate fractions larger than 10 µm diameter were collected at near daily intervals between early March and late May in 2018. Network analysis identified two major modules representing bacteria co-occurring with diatoms and with dinoflagellates, respectively. The diatom network module included known sulfate-reducing *Desulfobacterota* as well as potentially sulfur-oxidizing *Ectothiorhodospiraceae*. Metaproteome analyses confirmed presence of key enzymes involved in dissimilatory sulfate reduction, a process known to occur in sinking particles at greater depths. Our results indicate the presence of sufficiently anoxic niches in the particle fraction of an active phytoplankton bloom to sustain sulfate reduction, which may have implications for algal-bacterial interactions and carbon export during blooms.

## Introduction

Microalgae inject gigatons of organic carbon into coastal oceans every year (Field et al. 1998) and phytoplankton blooms represent primary productivity hotspots. It has been estimated that over 90% of algal-produced carbon is consumed by heterotrophic bacteria in the immediate vicinity of algal cells, the phycosphere, during typical bloom situations (Seymour et al. 2017). Some of these bacteria are directly associated with living algal cells, or with sinking aggregates of senescent or dead algae, and therefore play an important role in the biological carbon pump. Despite their importance in carbon sequestration (e.g., Bligh et al. 2022), vertical connectivity (Mestre et al. 2018), and their documented complexity (e.g. Reintjes et al. 2023), particle-associated (PA) bacterial communities are less well understood than their free-living counterparts. In fact, they are often overlooked due to the sometimes fragile nature of particles, in combination with the practice of pre-filtration to exclude larger organisms prior to molecular analyses (e.g. Simon et al. 2002, Thiele et al. 2015, Heins et al. 2021).

A defining feature of marine particles are the steep chemical and redox gradients that PA bacterial communities are exposed to, compared to free-living, planktonic bacteria. These gradients have been intensively studied in the context of bathypelagic sinking particles (i.e., marine snow), which harbor anaerobic micro-niches enabling, e.g., microbial sulfate reduction (Bianchi et al. 2018, Raven et al. 2021). In the photic zone, anaerobic micro-niches have been less widely investigated and the prevalence of sulfate-reducing bacteria is uncertain in the proximity of oxygen-producing living algal cells. Aggregates composed of living algae are rather known for a diverse microbiome of aerobic, heterotrophic bacteria that degrade complex algal organic matter, such as polysaccharide-rich exudates (Enke et al. 2019, Reintjes et al. 2023), although anaerobic metabolism such as diazotrophy is also known to occur (Riemann et al. 2022). Within this microbiome, bacteria are selected by factors such as host physiology and genotype (Ahern et al. 2021), the surrounding environment (Barreto Filho et al. 2021) and via stochastic processes (Stock et al. 2022). It is methodologically challenging to study associations between algal and bacterial taxa during natural phytoplankton blooms by direct observations, due to the transient nature of these associations (Heins et al. 2021, Seymour et al. 2017), as well as innate complexities and rapid dynamics of algal and bacterial communities during bloom events (Teeling et al. 2016). High-resolution temporal co-occurrence analysis, in combination with measurement of microbial functional potential, offers an indirect way to infer algae-bacteria associations, which can facilitate generating hypotheses about specific interactions and their potential functional implications.

Here, we investigated microbial community dynamics of PA bacterial and eukaryotic taxa during a spring phytoplankton bloom in the southern North Sea at the long-term ecological research site Helgoland Roads (Wiltshire 2010) in the year 2018. We aimed to identify PA bacterial taxa co-occurring with the major eukaryotic taxa (diatoms and dinoflagellates) during the bloom. We hypothesized that bacteria co-occurring with diatoms would be compositionally and functionally distinct from those co-occurring with dinoflagellates. Using 16S and 18S rRNA gene amplicon data from a well-resolved time series (near daily sampling) collected between early March and late May in 2018, we constructed co-occurrence networks focusing exclusively on bacteria-eukaryote co-occurrences in particles larger than 10 µm. In addition, we addressed bacterial functional gene expression by analysis of metaproteomes from three selected time points during the bloom.

## Materials and Methods

### Sampling and sample processing

A large volume of subsurface seawater (40 L - 140 L, 1 m depth) was sampled in the morning at near daily intervals between beginning of March and end of May in 2018 at the station long term ecological research (LTER) site Helgoland Roads (50° 11.3’ N, 7° 54.0’ E; DEIMS.iD: https://deims.org/1e96ef9b-0915-4661-849f-b3a72f5aa9b1) near Helgoland in the south-eastern North Sea. Secchi depth was measured from the vessel on site. For chlorophyll *a* (chl *a*) analysis, sample filtration was carried out in a laboratory under dim light to avoid the loss of pigments during the filtration process. We used a combined method of Zapata et al., (2000) and Garrido et al., (2003) for chl *a* extraction and analysis. Pigments were separated via high-performance liquid chromatography (HPLC) (Waters 2695 Separation Module), and detected with a Waters 996 Photodiode Array Detector. For 16S and 18S rRNA gene amplicon sequencing and metagenome analysis, plankton biomass from a 1 L seawater subsample was filtered using 10 µm pore size polycarbonate membrane filters (47 mm diameter, Millipore, Schwalbach, Germany) to separate PA microbes from smaller size fractions. At three selected time points during the bloom (Julian days 107, 128 and 144, representing early-, mid- and late bloom phases), a separate filtration was performed using larger (142 mm diameter, Millipore) 10 µm polycarbonate membrane filters for metaproteomic analysis. In order to maximize biomass harvest while avoiding clogging, filtered volumes varied between 15 and 30.5 L per filter for metaproteomic analysis.

### rRNA gene amplicon sequencing

Samples from 52 time points throughout the bloom were collected and analyzed. DNA was extracted from the filters using the Qiagen DNeasy Power soil Pro kit (Qiagen, Hilden, Germany) according to the manufacturer’s instructions. Dislocation of microbial cells from the filters and mechanical lysis were achieved by bead beating in a FastPrep 24 5G (MP Biomedicals, Irvine, CA, USA). DNA concentrations were measured at a Qubit 3.0 fluorometer (Invitrogen, Carlsbad, CA, USA). Extracted DNA was amplified with primer pairs targeting the V4 region of the 16S rRNA gene [515f: 5‘-GTGYCAGCMGCCGCGGTAA-3‘, 806r: 5‘-GGACTACNVGGGTWTCTAAT-3’ (Walters et al., 2016)] and the V7 region of the 18S rRNA gene [F-1183mod: 5’-AATTTGACTCAACRCGGG-3’, R-1443mod: 5’- GRGCATCACAGACCTG-3’] (Ray et al. 2016) coupled to custom adaptor-barcode constructs. PCR amplification and Illumina MiSeq (Illumina, San Diego, CA, USA) library preparation and sequencing (V3 chemistry) were carried out by LGC Genomics (LGC Genomics, Berlin, Germany).

Sequence reads free of adaptor and primer sequence remains were processed using the DADA2 package (v1.2.0) in R (Callahan et al., 2016). In summary, forward and reverse Illumina MiSeq reads were truncated to 200 bp, filtered (maxEE = 2, truncQ = 2, minLen = 175), dereplicated and error rates were estimated using the maximum possible error estimate from the data as initial guess. Sample sequences were inferred, paired forward and reverse reads were merged and chimeric sequences were removed using the removeBimeraDenovo function. The resulting amplicon sequence variants (ASVs) were taxonomically classified using the Silva database (nr 99 v 138.1) for 16S rRNA and the PR2 database (version 4.13, minboot: 50) for 18S rRNA sequences using the build-in RDP classifier. 16S rRNA gene amplicon reads classified as chloroplasts and mitochondria, as well as the 18S rRNA gene reads classified as *Metazoa* (zooplankton) were removed prior to downstream analyses (a single time point was excluded due to suspected contamination).

Co-occurrence networks were generated in R using Spearman rank correlation, as described previously (Bengtsson et al. 2017, correlation coefficient >0.7, adjusted p<0.01). We considered exclusively correlations between 18S and 16S ASVs and the final network was plotted using the igraph R package.

### Construction of a metagenome-based database

Illumina sequencing was performed at the Max Planck Genome Centre Cologne on DNA extracted from both 3-10 µm- and >10 µm filter fractions from eight selected time points (19 March, 12 April, 17 April, 26 April, 08 May, 11 May, 22 May and 29 May 2018 - Julian days 78, 102, 107, 116, 128, 131, 142 and 149), using the HiSeq 2500 platform and 2 x 150 bp chemistry. Filtration and trimming of the reads were done as described previously (Francis et al. 2021). Read quality for each sample was confirmed using FastQC v0.11.9 (Andrews 2014). Assembly was performed with MEGAHIT v1.2.9 (Li et al. 2015) (kmer length: 21). Contigs below 2.5 kbp were removed using anvi-script-reformat-fasta from within Anvi’o v6.2 (Eren et al. 2015). Genes were predicted using Prodigal v2.6.3 (Hyatt et al. 2010) as implemented in the Prokka v1.11 annotation pipeline (Seemann 2014). These genes sequences were subsequently combined with a list of common contaminants to generate a database for peptide-spectrum-matching (PSM) after elimination of redundant sequences (97% redundancy) with CD-Hit (Fu et al. 2012, Li and Godzik 2006).

### Metaproteomics

Proteins were extracted from biomass filters of the 17th of April, 8th of May and 24th of May 2018 (Julian days 107, 128 and 144) as described previously (Schultz et al. 2020, Schultz 2022) and analyzed with liquid chromatography-tandem mass spectrometry in triplicates. Briefly, proteins were extracted via bead-beating followed by acetone precipitation. The extracts were separated and fractionated by 1D SDS-PAGE and in-gel trypsin digested. After desalting and concentration of the peptides using C18 Millipore® ZipTip columns, the samples were measured with an Orbitrap Velos^TM^ mass spectrometer (ThermoFisher Scientific, Waltham, MA, USA). After conversion into mgf file format using MS convert in ProteoWizard (Palo Alto, CA, USA), spectra were matched against a metagenome-based database containing 14,764,755 entries. Mascot (Matrix Science, London, UK) and Scaffold (Proteome Software, Portland, OR, US) were used for peptide-spectrum-matching, protein identification and protein grouping. Instead of setting an FDR-threshold, identification of protein groups was based on number of peptide-matches with a minimum of two, a protein threshold of 99% and a peptide threshold of 95%. Identified protein groups were annotated via Prophane v6.2.3 (Schiebenhoefer et al. 2020), using the Uniprot-TrEMBL (as of September 2021) and NCBI nr (as of February 2022) databases for taxonomic and EggNog v5.0.2 and Tigrfams 15 databases for functional annotation with Prophane default settings. Taxonomic information for *Desulfobacterota* proteins was manually confirmed against the most recent ncbi nr database (blastp, https://blast.ncbi.nlm.nih.gov, September 2023). TrEMBL taxonomic information for *Desulfobacterales* was manually curated and set to phylum level for better comparability with the Silva database. Relative abundances were calculated as Normalized Spectral Abundance Factor (NSAF) values using the quantification method “max_nsaf” integrated in Prophane (Schiebenhoefer et al., 2020). Briefly, SAF values were calculated by dividing exclusive unique spectrum counts for each protein group by protein length of the longest sequence in that protein group. For normalization, SAF values were then divided by the sum of all SAF values of the sample.

### Data visualization

Stacked bar plots for rRNA gene amplicon data and metaproteomics were created with R version 4.3.0 using the tidyverse package (Wickham et al. 2019) in combination with the svglite, polychrome, patchwork, glue and ggnested packages.

## Results

### Particle microbial community dynamics during the course of the bloom

The spring phytoplankton bloom in 2018 was characterized by an initial dominance of diatoms (*Bacillariophyceae*), followed by an increase in dinoflagellate (*Dinophyceae*) relative abundances after the bloom peak (peak in chl *a* concentration, Fig. 1a and b). The algal bloom (chl *a* measurements) peaked around the 26th of April (Julian day 116), coinciding with a decrease in water clarity, as indicated by Secchi depth rising at the onset of the bloom. 16S rRNA gene amplicon sequencing revealed a total of 17615 ASVs in the particle bacterial microbiomes, which consisted mainly of *Proteobacteria (*e.g. *Spongibacteriaceae, Methylophagaceae, Rhodobacteriaceae)*, *Bacteroidetes* (e.g. *Polaribacter*), *Verrucomicrobia* (e.g. *Persicirhabdus*) and *Planctomycetes* (e.g. *Phycisphaeraceae*). Further, *Actinobacteria* (e.g. *Illumatobacter*), and, notably, *Desulfobacterota* were also relatively abundant (Fig. 1c).

**Figure 1:**
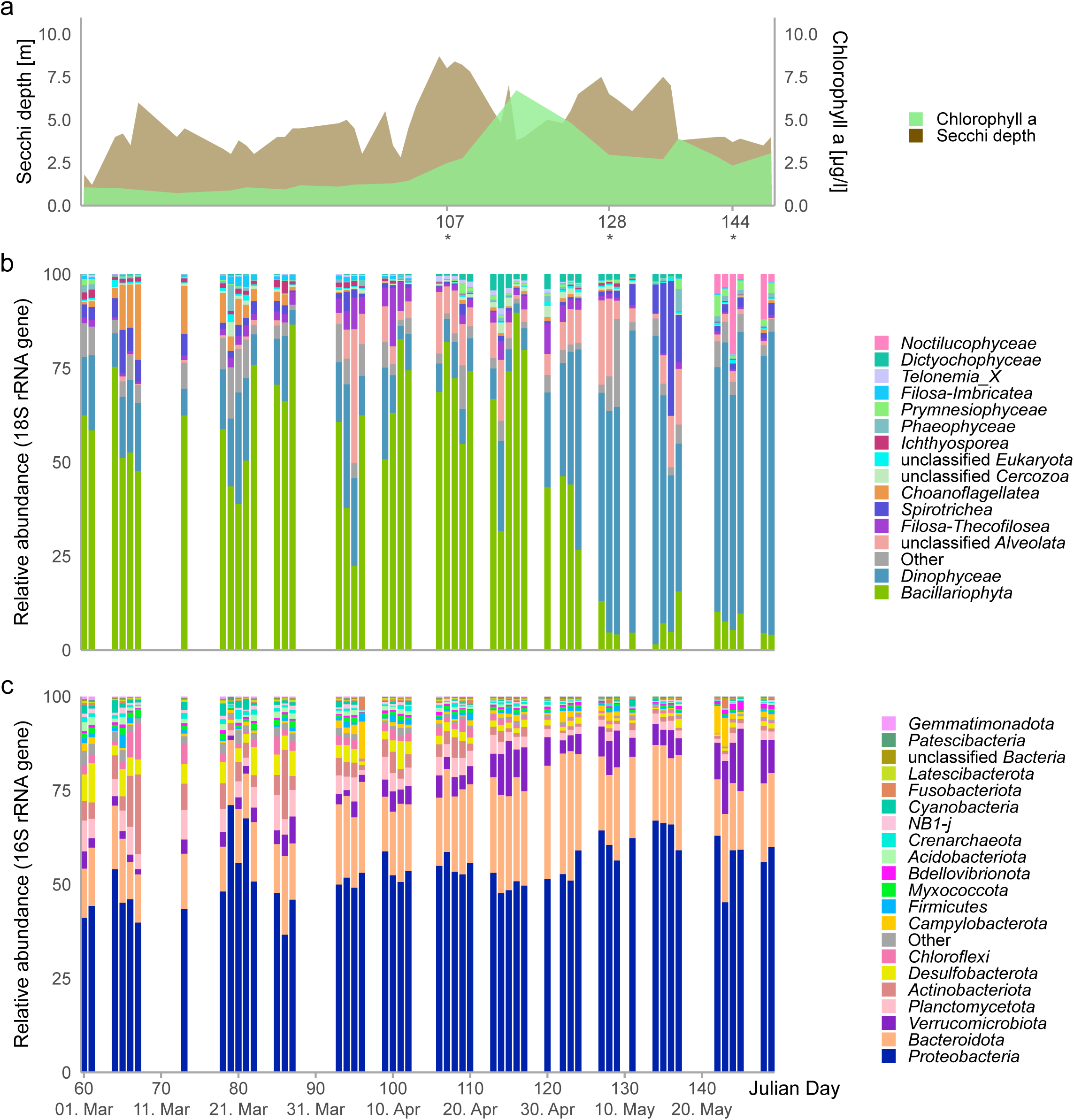
Progression of the 2018 spring bloom by Helgoland, North Sea. (a) Chl *a* (green) peaked around the 26th of April (Julian day 116), coinciding with a drop in Secchi depth (brown). Asterisks indicate time points (Julian days) for which metaproteomic sampling was performed. (b) 18S rRNA gene amplicon sequencing revealed that diatoms (*Bacillariophyta*) were the most abundant phytoplankton lineage, although dinoflagellates (*Dinophyceae*) dominated after the chl *a* peak. (c) 16S rRNA gene amplicon sequencing showed the highest relative abundances of *Desulfobacterota* (yellow) during the first half of the bloom.

### Taxonomic and functional composition as detected by metaproteome analysis

Metaproteome sampling was performed on three time points that correspond to the early, mid and late phase of the bloom (Julian days 107, 128 and 144), as indicated in Fig. 1a). We detected 4,584 protein groups in >10 µm filter fractions from these sampled time points. Metaproteomic analyses confirmed high abundances of *Proteobacteria* and *Bacteroidetes* in the bacterial fraction (Fig. 2a), and dominance of *Bacillariopyhta* in the eukaryote fraction (Fig. 2b), with eukaryote proteins making up 90% of all proteins at the first selected time point. The proportion of eukaryotic proteins was reduced to 70% in the last selected time point (144), while bacterial proteins became comparably more abundant (25% on day 144 compared to 4% on day 107). Functional analysis revealed a predominance of eukaryotic proteins involved in metabolism, including energy production and conversion, and a shift towards expression of proteins relevant in cellular processes and signaling over the course of the bloom (Fig. 2c).

**Figure 2:**
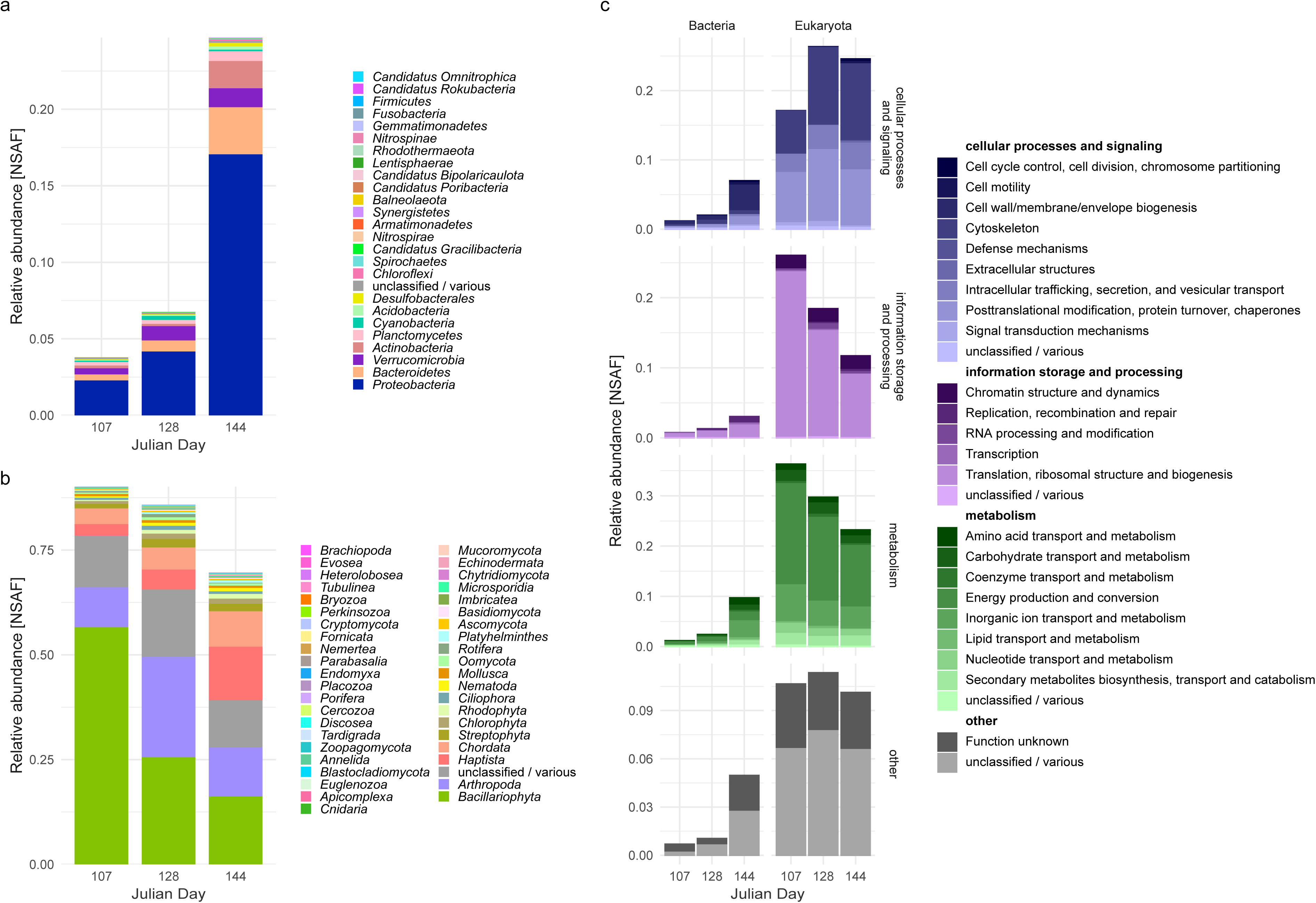
Taxonomic and functional annotation of particle metaproteomes from three selected time points during the bloom. (a) The proportion of bacterial proteins increased during the course of the bloom (b) Eukaryotic proteins made up the majority of identified proteins and showed an initial strong dominance of *Bacillariophyta* (c). Functional annotation of metaproteomes indicated a high contribution of metabolism-related eukaryotic proteins early in the bloom, while cellular processes and signaling increased during later time points.

### Co-occurrence network analysis

Co-occurrence analysis resulted in distinct network modules centered around diatom- and dinoflagellate 18S rRNA gene amplicon sequence variants (ASVs). Based on significant positive correlations (Spearman Rho>0.7, corrected p>0.01) between eukaryotic 18S rRNA gene and prokaryotic 16S rRNA gene ASVs, including the 51 analyzed time points during the 2018 spring bloom, the network was dominated by three major distinct modules (Fig. 3a). Two of these modules were dominated by eukaryotic ASVs belonging to diatoms and dinoflagellates, respectively, while the third module mostly contained diatom ASVs and was linked to the dinoflagellate-dominated network module. Along the time line of the bloom, these modules roughly corresponded to the phytoplankton taxa prevalent in the early stages of the bloom (module I, mainly diatoms), during the late stages of the bloom (module II, mainly dinoflagellates) and mid-bloom around peak chl *a* (module III, Fig. 1). The composition of bacterial ASVs that co-occurred with diatoms and dinoflagellates is depicted in Fig. 3b. Eleven ASVs belonging to the *Desulfobacterota* were part of the main diatom- dominated network module I and co-occurred exclusively with diatoms, including the genus *Desulfosarcina*, and the families *Desulfocapsaceae* and *Desulfobulbaceae* as well as the lineage Sva1033 (Ravenschlag et al. 1999). In addition, 4 ASVs classified as *Ectothiorhodospiraceae* (genus *Thiogranum*) also co-occurred with diatoms (Fig. 3b).

**Figure 3:**
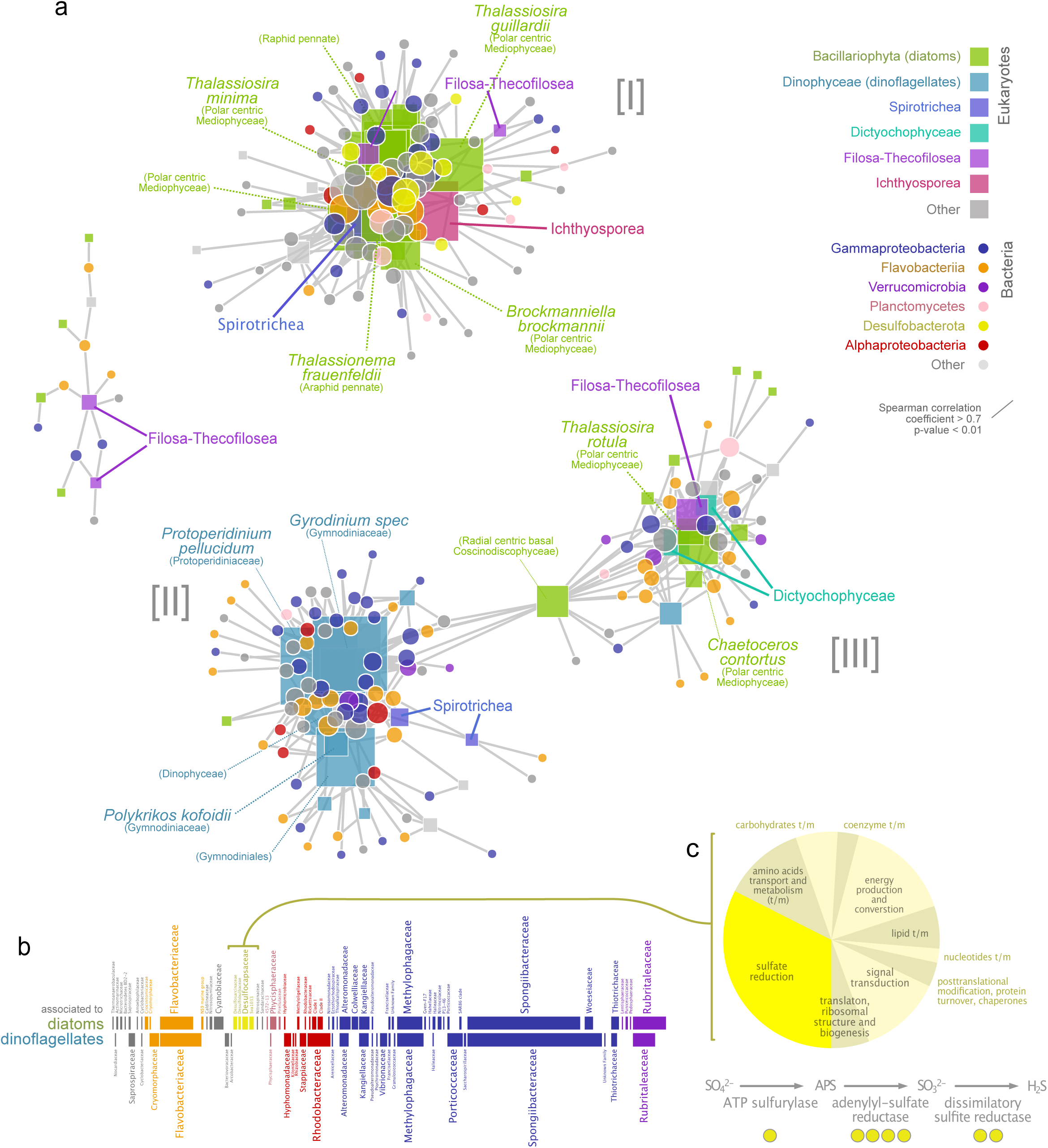
**Co-occurrence of eukaryotes and bacteria, and detected protein groups of *Desulfobacterota*** (a) A network analysis of 18S rRNA (triangles) and 16S rRNA (circles) gene ASV co-occurrences resulted in two major network modules containing diatom [I] and dinoflagellate [II] 18S ASVs, respectively, as well as one mixed module [III]. The network was calculated based on Spearman correlations (r >0.7, p < 0.01) exclusively between 18S ASVs and 16S ASVs. *Desulfobacterota* (yellow circles) were only associated with the diatom-dominated module I. The sizes of the symbols correspond to the number of significant correlations of the nodes (degree). (b) The bacterial taxa co-occurring with diatoms and dinoflagellates, respectively, are displayed as horizontal bars with a length proportional to the number of ASVs belonging to each lineage. *Desulfobacterota* (yellow) as well as potentially sulfur-oxidizing *Ectothiorodospiraceae* (*Gammaproteobacteria*, blue) were positively associated with diatoms but not with dinoflagellates. (c) The pie chart displays relative abundances for all 19 protein groups classified as belonging to *Desulfobacterota* during all three time points sampled for metaproteomics. Of these, seven protein groups (32%, bright yellow) represented enzymes involved in dissimilatory sulfate reduction. Yellow circles indicate which of the key enzymes in this metabolic pathway were detected.

### Analysis of Desulfobacterota proteins

Out of the 4,584 protein groups detected by metaproteome analysis, 19 were classified as belonging to different *Desulfobacterota* orders (Table S1). The relative abundance of predicted *Desulfobacterota* proteins averaged over all time points in the metaproteome was 0.14% (expressed as normalized spectral abundance factor - NSAF). This order of magnitude corresponded well to the relative abundance of *Desulfobacterota* ASVs in the microbiome at the same time points (average 0.38%, Fig 1c). Remarkably, no less than 32% of the *Desulfobacterota* metaproteome fraction (in terms of protein abundance) consisted of key enzymes for dissimilatory sulfate reduction (Fig. 3c). Of these, ATP sulfurylase, both the alpha and the beta subunits of dissimilatory sulfite reductase (DSR) and adenylyl-sulfate (adenosine-5’-phosphosulfate) reductase (APS reductase) were detected. We used a conservative threshold for protein identification of at least two matching peptides to consider a protein as validly detected.

## Discussion

The succession we observed during the 2018 spring bloom, with an initial dominance of diatoms, followed by dinoflagellates, using 18S rRNA gene amplicon sequencing agrees with the typical phytoplankton community succession at Helgoland Roads (Wiltshire et al. 2008, Käse et al. 2020). Metaproteome analysis highlighted the dominance of eukaryotic proteins in the sampled particles, making up 70-80% of detected proteins, most of which belonged to the major diatom phytoplankton. The high prevalence of proteins involved in basic metabolism such as energy production and conversion is consistent with the sampled >10 µm particle fraction mostly comprising of living phytoplankton cells, especially at the beginning of the bloom. However, bacterial proteins were present in all of the three selected time points, with a notable peak at day 144, and their taxonomic composition was very similar to that observed via 16S rRNA gene amplicon sequencing. Overall, the composition of particle bacterial microbiomes agreed with other reports from similar environments (Reintjes et al. 2023, Schultz et al. 2020, Heins et al. 2021, Crump et al. 1999).

As hypothesized, bacteria co-occurring with abundant diatoms formed a distinct network module, with taxa that were different from those co-occurring with dinoflagellates in a second distinct module. A third network module contained mostly diatom taxa, but also one dinoflagellate ASV and other phytoplankton taxa. These network patterns offer an alternative way to visualize the temporal dynamics of the bloom, and should not be interpreted as evidence of physical interactions between taxa (Röttjers & Faust 2018). However, one pattern that is striking in our network analysis is the exclusive co-occurrence of *Desulfobacterota* with diatoms.

We observed a high relative abundance of *Desulfobacterota* (in total 0.38% of bacterial amplicon reads) in particles (>10 µm), especially in the early phase of the bloom when diatoms were dominating. Despite the limited resolution of metaproteome data compared to DNA-based methods, desulfobacterial proteins were represented at all selected time points (in total 0.14% of metaproteomes). Network analysis further highlighted the temporal co- occurrence of *Desulfobacterota* with several diatom taxa. This raises the question of the niche filled by these anaerobic bacteria during an active phytoplankton bloom.

Diatoms, such as *Thallassiosira* spp. and *Thalassionema* spp., which were co-occurring with *Desulfobacterota* in this study (Fig. 3a), are known to form aggregates (Thornton 2002). For example, several species of *Thalassiosira* extrude long chitin fibrils, which prevent sinking and bind exopolymeric substances (EPS) also produced by the algal cells (Den et al. 2023, Herth & Barthlott 1979). This creates a favorable environment for bacteria to attach, which can in turn stimulate algal EPS production (Gärdes et al. 2011). EPS makes particles adhesive and thus bacteria can be captured in this sticky EPS layer. Smaller particles can aggregate to larger particles by collision and adhesion, in particular during phytoplankton blooms with high particle densities. Such aggregates can feature high numbers of living, photosynthesizing algal cells (Thornton 2002) which produce ample oxygen during the day, but at night respiration by the algal cells and their surrounding bacteria may deplete oxygen sufficiently for low oxygen or even anaerobic micro-niches to form.

*Desulfobacterota* have not been identified as frequent members of diatom microbiomes in either cultures or in the field so far (Helliwell et al. 2022). However, a recent global survey of the diatom interactome detected positive correlations between diatoms and sulfate- reducing bacteria in samples from the Tara Oceans expedition (Vincent & Bowler 2020). In addition, *Desulfobacterota* have previously been repeatedly detected in particle-associated communities in the photic zone (Crump et al. 1999, Liu et al. 2020, Hallstrøm et al. 2022). *Ectothiorhodospiraceae* are purple sulfur bacteria (belonging to *Gammaproteobacteria*), that are anaerobic to microaerophilic (Imhoff 2022). They can oxidize H_2_S, e.g., produced via dissimilatory sulfate reduction by members of the *Desulfobacterota*. Our observation of *Desulfobacterota* as well as *Ectothiorhodospiraceae* co-occurring with diatoms is consistent with potential sulfur cycling during the diatom-dominated phase of this phytoplankton bloom under anoxic to very low oxygen conditions. While co-occurrence of diatom and *Desulfobacterota* rRNA genes does not by itself indicate that sulfate reduction is taking place in diatom-derived particles, detection of key functional enzymes by metaproteomics suggests that *Desulfobacterota* were actively carrying out sulfate reduction and thus gaining energy through anaerobic respiration during the bloom. The higher detection of these enzymes especially at the last time point (Julian day 144, Table S1) can likely be attributed to the larger proportion of bacterial proteins at this time point, when the diatom bloom was declining.

Temporal associations, such as detected by our co-occurrence network analyses, have to be interpreted with caution as additional variable factors that were not taken into account may influence the observed correlations. Importantly, we cannot quantitatively assess the influence of underlying sediment microbiomes at the rather shallow sampling site (6-10 m depth depending on tide), which frequently become resuspended during times of heavy wind thereby introducing anaerobic microbes to the pelagic environment. Although we did not find a clear link between Secchi depth, wind conditions and *Desulfobacterota* relative abundances in this study (Fig. 1, Fig. S1), the detected ASVs (e.g. classifying as *Desulfosarcina*, *Desulfocapsaceae*, *Desulfobulbaceae*) are closely related to typical benthic lineages (Ravenschlag et al. 1999), suggesting that they originated from the underlying sediments. The diatoms co-occurring with *Desulfobacteria* were primarily classified as common pelagic genera such as *Thalassiosira* and *Thalassionema*. However, the genus *Brockmanniella*, which can also inhabit benthic biofilms (Hernández Fariñas et al. 2017), was also found within network module I, suggesting some influence of sediment resuspension. In shallow environments, benthic and pelagic microbiomes are in close contact within the photic zone, and seeding of pelagic particles by benthic microbes (and *vice versa*) is likely frequent, which may explain the observed co-occurrences. However, our methodology does not allow us to determine the physical proximity of *Desulfobacterota* and diatom cells. The observed co-occurrences may therefore reflect a purely temporal association between pelagic diatoms and resuspended sediment bacteria. Indeed, demonstrating a potential physical association between *Desulfobacterota* and diatoms would require *in situ* microscopic investigation using for example taxon- or gene- specific fluorescent probes. Likewise, it has yet to be clarified, whether such temporal or physical associations are annually recurrent, under what specific conditions they occur, and which specific diatom taxa are involved. Thus, further studies are needed to confirm whether our results are representative for phytoplankton blooms in shallow water coastal areas.

Our results highlight the complexity of particle microbiomes and corroborate the need to study algae-bacteria particles as spatially heterogeneous entities. The microbiomes of aggregate-forming phytoplankton may indeed be similarly complex as those of animals and other multicellular organisms, featuring distinct micro-niches with sub-microbiomes (analogous to e.g., human skin vs. gut microbiomes), whose compositions depend on the chemical environment on a small spatial scale. Anaerobic micro-niches in phytoplankton- derived aggregates may affect carbon cycling insofar, as sulfate reduction can chemically alter particulate organic matter, rendering it resistant to microbial degradation (Raven et al. 2021). Anaerobic niches in marine phytoplankton-derived particles were already frequently reported in the context of nitrogen fixation (i.e., diazotrophy, e.g. Riemann et al. 2022).

Recent work has shown that many *Desulfobacterota* are in fact capable of both diazotrophy and sulfate reduction (Liesirova et al. 2023) and that diazotrophic *Desulfobacterota* can move towards phytoplankton-derived organic matter via chemotaxis. Similar to diazotrophs, sulfate reducers in turn provide additional niches for bacteria which scavenge their metabolic products, i.e., sulfur oxidizers such as the observed *Ectothiorhodospiraceae*, leading to higher metabolic diversity and complexity within particles. Considering the significance of marine particle microbiomes in carbon sequestration, it is vital to understand the consequences of the metabolic and compositional complexity in particle microbiomes, and the resulting microbial interactions.

## Author Contributions

RS conducted the metaproteomic analyses together with DS and DZ, analyzed the data, prepared figures and contributed to writing the manuscript. MSF performed amplicon sequencing analyses together with AB. FW prepared metagenomics databases together with HT and KJH. PAM performed mass-spectrometry measurements together with DB. JB contributed to data analysis and figure preparation. IVK contributed chl *a* and Secchi depth data together with KHW. TU contributed to data analysis and interpretation. KR supervised metaproteome analysis and conceived the study together with MMB. MMB supervised and contributed to amplicon data collection, analysis, interpretation and wrote the manuscript with significant input from RS, HT, TU and IVK. All authors approved the final version of the manuscript.

## Funding

This study was funded by the Deutsche Forschungsgemeinschaft (DFG) in the framework of the research unit FOR2406 ‘Proteogenomics of Marine Polysaccharide Utilization (POMPU)’ by grants of KR/MMB (RI 969/9-2), DB (BE 3869/4-3) and HT (TE 813/2-3).

Access to Alfred-Wegener-Institut Helmholtz-Zentrum für Polar- und Meeresforschung (AWI) infrastructure was ensured through the Helmholtz Association’s LK-II performance category program (AWI_BAH_o 1).

## Acknowledgments

We thank the Biological Station Helgoland, Alfred-Wegener-Institut Helmholtz-Zentrum für Polar- und Meeresforschung and the crews from FS Aade and FS Uthörn for help with sampling, analyses, logistics, and providing lab space. Jan Rockstroh and Kira Wisnewski assisted in sample processing for metaproteomics. Thomas Ben Francis assisted in metagenome database construction. We are also grateful to the whole POMPU consortium for helpful discussions, especially Rudi Amann, Jens Harder, Saskia Kalenborn and Jan-Hendrik Hehemann.

## Data Availability Statement

Metagenome raw reads and assembled contigs used for metaproteome assignment are publicly available at the European nucleotide archive under the project accession number PRJEB38290. 16S- and 18S-rRNA gene amplicon data are also available via ENA under the project accession number PRJEB51816. Mass spectrometry proteomics data have been deposited to the ProteomeXchange Consortium via the PRIDE (Perez-Riverol et al., 2022) repository with dataset identifier PXD035982. Original mass spectrometry proteomics data can be accessed for review (**Username:** **Password:** umim4qCW) and will be made publicly available upon manuscript acceptance.

