## Supplementary material for "Marine particle microbiomes during a spring diatom bloom contain active sulfate-reducing bacteria"

Supplementary figures	2

Supplementary tables	4

### Supplementary figures

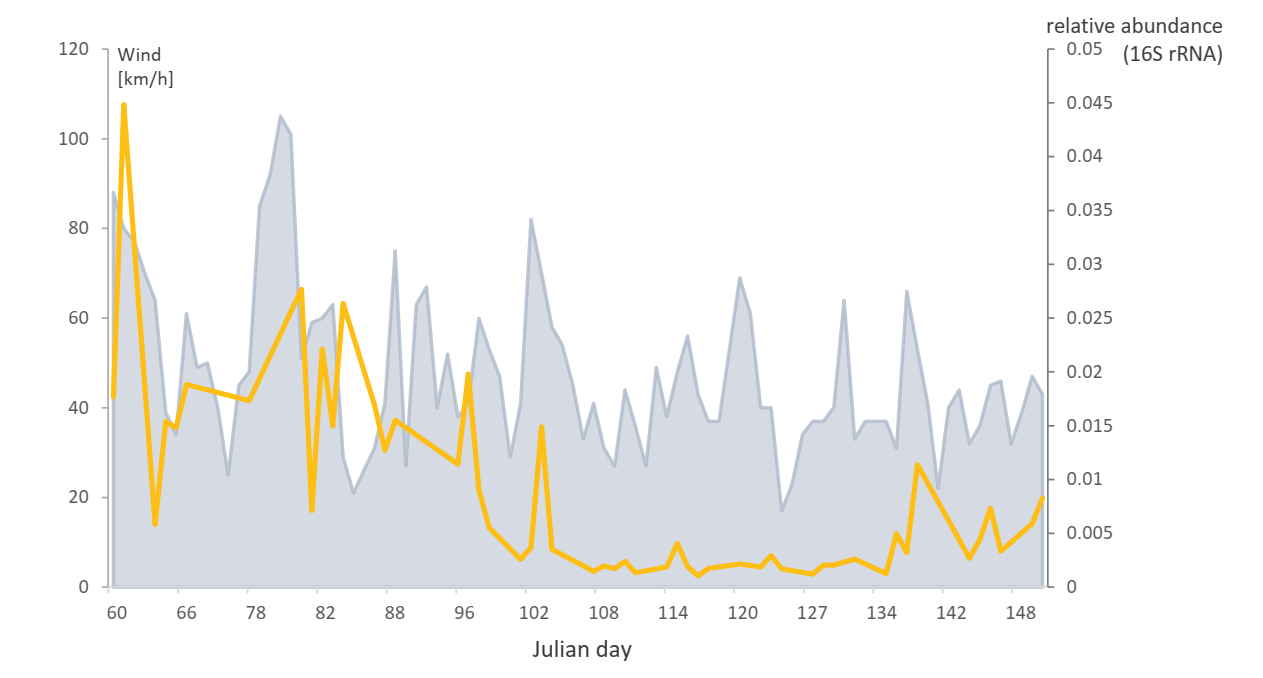

**Supplementary Figure 1:** Relative abundance of *Desulfobacterota* in relation to wind conditions during the 2018 spring bloom. Relative abundances of *Desulfobacterota* are based on 16S rRNA gene amplicon ASVs. Wind maxima are based on DWD Helgoland station via wetterkontor.de.

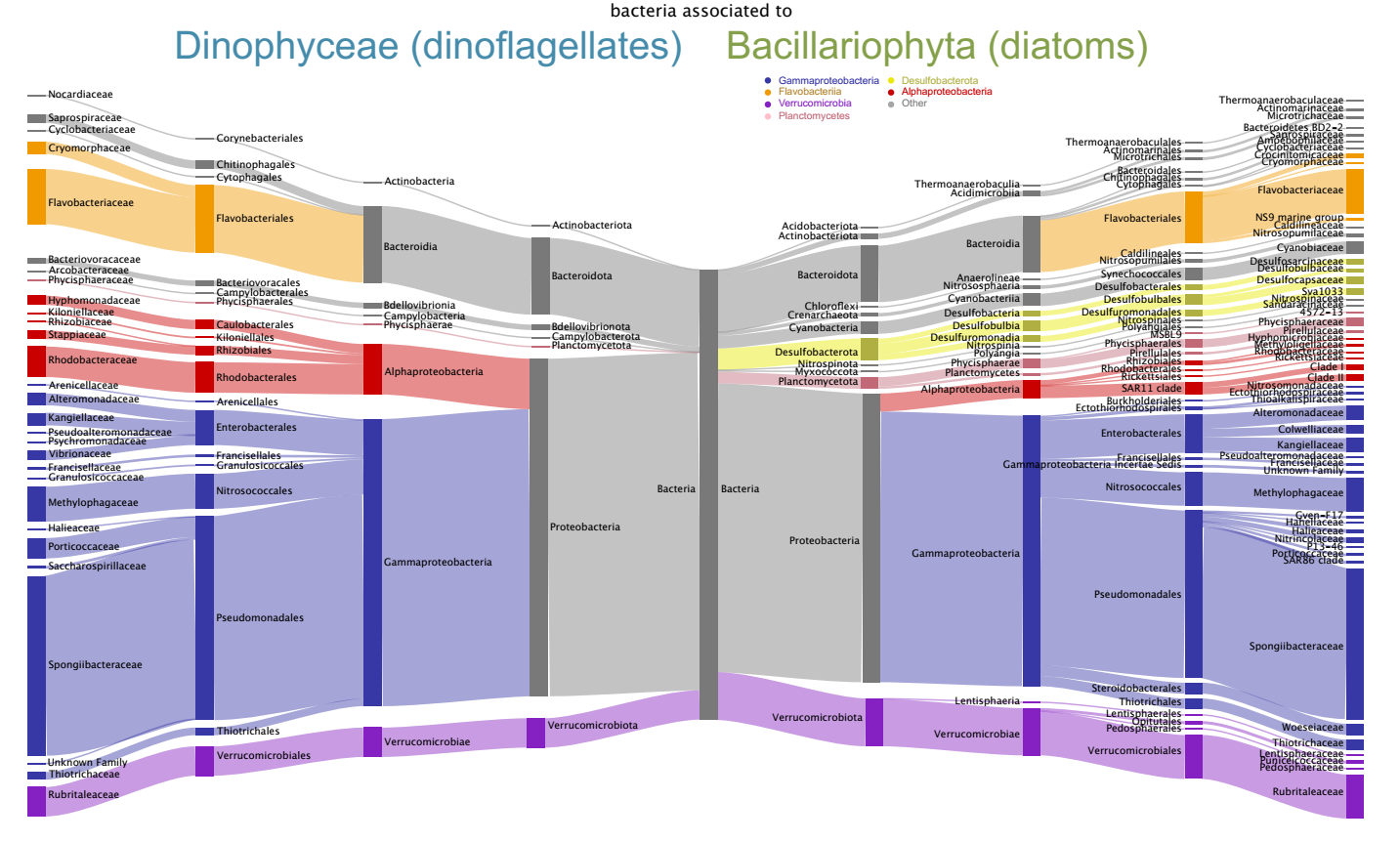

**Supplementary Figure 2:** Association of *Desulfobacterota* and *Ectothiorodospiraceae* with diatoms. Distribution of bacterial ASVs based on 16S rRNA gene amplicon data. Temporal associations to diatoms and dinoflagellates were taxonomically distinct.

### Supplementary tables

| #pg | Protein function | NSAF | | | Tax_summary |
| --- | --- | --- | --- | --- | --- |
|  |  | Julian Day 107 | Julian Day 128 | Julian Day 144 |  |
| 112 |  | 0 | 0 | 0.00016602 | *Desulfobulbaceae* sp. |
| 182 | Adenylylsulfate reductase | 0 | 0 | 0.00015253 | *Desulfobulbaceae* sp. |
| 293 |  | 5.7524E-05 | 7.7272E-05 | 0 | *Desulfosarcina* sp. |
| 411 | Adenylylsulfate reductase | 0 | 0 | 0.00025035 | *Desulfosarcina* sp. |
| 440 | Dissimilatory sulfite reductase | 0 | 0 | 0.00013727 | *Desulfobacteraceae* sp. |
| 540 |  | 0.00023951 | 0 | 0 | *Desulfobulbaceae* sp. |
| 576 | Dissimilatory sulfite reductase | 0 | 0 | 0.00018468 | *Desulfobulbaceae sp.* |
| 666 |  | 0 | 0 | 0.00015508 | *Desulfosarcina* sp. |
| 743 |  | 5.4294E-05 | 1.8454E-05 | 8.2017E-05 | *Desulfobacterales* sp. |
| 810 |  | 0.0002416 | 0.00015172 | 0.00011497 | *Desulfobacterales* sp. |
| 954 | ATP sulfurylase | 0 | 0 | 0.00016176 | *Desulfobulbaceae* sp. |
| 960 |  | 0 | 0.00017834 | 0.00034704 | *Desulfobacterales* sp. |
| 1214 |  | 0 | 0 | 8.941E-05 | *Desulfobacterales* sp. |
| 1321 |  | 0 | 0.00010197 | 0.00015095 | *Desulfosarcina* sp. |
| 1513 |  | 0 | 0 | 0.00011631 | *Desulfofustis* |
| 2117 |  | 0 | 0 | 0.00026687 | *Desulfobacterales* sp. |
| 2147 | Adenylylsulfate reductase | 0 | 0 | 0.00037822 | *Desulfopila* sp. |
| 2745 |  | 0 | 0 | 0.00026719 | *Desulfofustis* |
| 2868 | Adenylylsulfate reductase | 0 | 0 | 0.00011735 | *Desulfofustis* |

**Supplementary Table 1:** Seven key enzymes associated with the sulfur cycle detected with metaproteomics. ATP sulfurylase, adenylylsulfate-reductase and dissimilatory sulfite reductase assigned to five *Desulfobacterota* species could be detected. Functional annotation was based on EggNog v5.0.2, and the assignment of taxonomy was based on Trembl (as of Sept. 2021), and NCBI nr (as of Feb. 2022) database annnotations and manually confirmed against the most recent NCBI nr database via <https://blast.ncbi.nlm.nih.gov/> (blastp, Sept. 2023).
